# Multiphase coexistence capacity in complex fluids

**DOI:** 10.1101/2022.10.19.512909

**Authors:** Krishna Shrinivas, Michael P. Brenner

## Abstract

Complex fluids like the cytoplasm comprise hundreds of species organized into many coexisting phases. How molecular interactions, reflecting sequence, design, or functional constraints, dictate multiphase coexistence is a major open question. To answer this, we consider models of multicomponent fluids with both designed and random interactions. When crosstalk is introduced, we show that coexisting phases lose specificity beyond a common threshold. In a sequence model, we demonstrate that phase capacity is limited by sequence length and grows logarithmically/linearly with sequence length/number. These results provide a general route to program multiphase coexistence from molecular features.

## Introduction

Life is rife with spatially inhomogeneous materials. A prominent example is the cellular milieu which is organized into dozens of coexisting compartments called condensates [1,2]. Individual condensates contain tens of protein and nucleic acid components [1–4] whose interaction networks span a wide range of strengths and specificities [5–8] and often assemble by phase separation. A major outstanding question lies in predicting how microscopic interactions and molecular features, reflecting sequence-derived, functional, and evolutionary constraints, encode for macroscopic multiphase coexistence.

The conditions for thermodynamic equilibria and multiphase coexistence are particularly challenging to delineate in materials with many components. This has limited the characterization of phase behavior to fluids with very few components (<5) [9–11]. Rather than focus on exhaustive enumeration, an alternative strategy is to relate *statistical* ensembles of interacting molecules to their emergent properties. Motivated by pioneering work by Sear and Cuesta [12] and others [13], a number of mathematical approaches have emerged to dissect the point of the spinodal manifold i.e. concentrations where a fluid begins to spontaneously phase separate, for fluids with random and partially structured interactions [14–16]. These methods are limited to identifying the spinodal, without predicting the number or composition of coexisting phases at steady state. More recently, approaches from non-equilibrium thermodynamics have begun to make progress in tackling this question [17,18]. Phase-field models have been deployed to predict not only steadystate properties but also out-of-equilibrium dynamical behavior for randomly interacting fluids [17]. Numerical schemes rooted in linearly irreversible and mean-field thermodynamics [18] identify *potential* thermodynamic equilibria across a broad range of interaction parameters. Whether these distinct methods find similar steady-states is not understood although there is broad agreement when interactions are purely random. While characterizing random fluids represents a first step [17], interactions amongst molecules in cells and other complex fluids are non-random, reflecting correlations derived from sequence, functional, or evolutionary constraints. How non-random biomolecular interactions encode for multiphase coexistence is not known. This, in turn, limits our ability to design molecular features to program or engineer target macroscopic behavior.

In this paper, we investigate and characterize multiphase coexistence in fluids whose component interactions are non-random by combining phase-field simulations, mean-field thermodynamics, and eigen-mode counting arguments. We first explore a simple model in which components interact through strong, specific interactions within a group (or block) and through non-specific random crosstalk with all other components. When crosstalk is higher than a sharp threshold, we identify a transition where number and composition of coexisting phases change from those encoded by highly structured blocks to random fluids i.e., many decoherent phases. Inspired by interacting DNA sequences and low-complexity proteins, we then study a model where interactions and concomitant crosstalk are encoded by sequence-features, parametrized by length *R* and interaction strength *J*. With *N* randomly generated sequences, we find a maximum number (or capacity) of coexisting phases 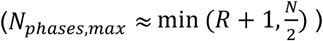 when interactions are strong. With weak or moderate interactions, we identify a universal scaling law connecting sequence features to multiphase capacity (*N_ph_* ∝ *N*log (*R*)). Together, our results provide an avenue to program multiphase coexistence by tuning molecular features.

## Model

We employ a regular solution or Flory-Huggins like formalism to describe the free-energy for a mixture of N interacting species in an inert solvent at fixed volume and temperature (SI Theory). The component volume fractions are defined as *ϕ_i_*. For simplicity, we assume equimolar fluids where interactions *ϵ_ij_* are purely heterotypic (i.e., no self-interactions) and derived from an underlying model of interactions (Figure 1).

**Figure 1:**
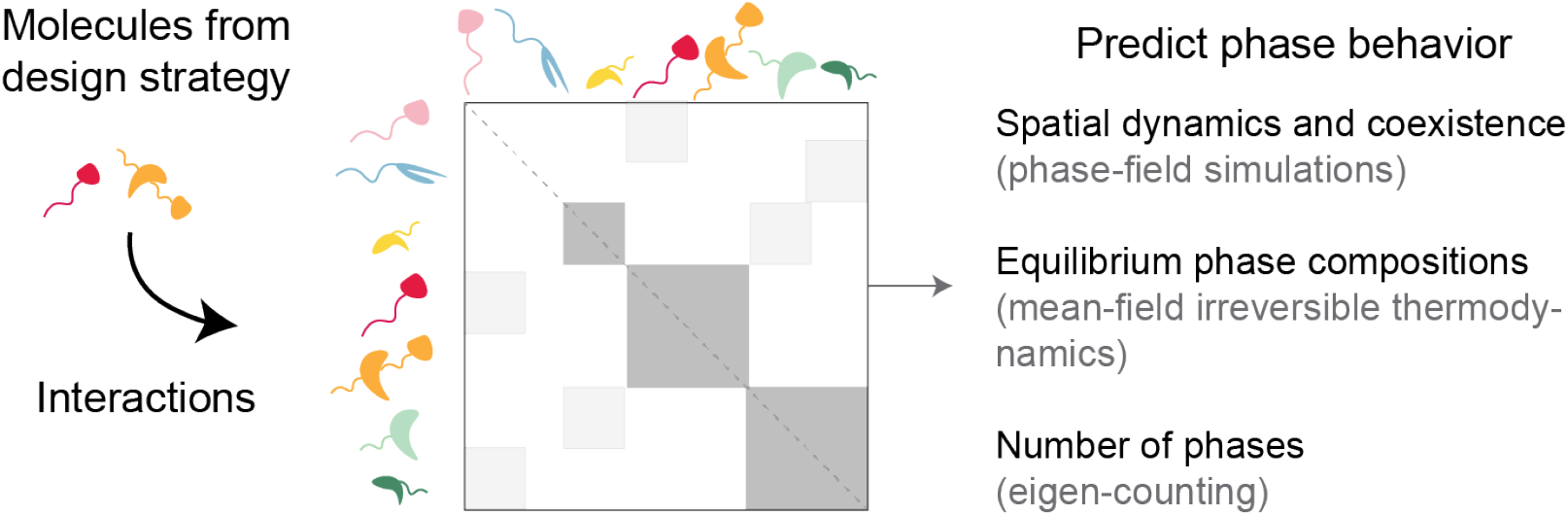
Model for probing multiphase coexistence in multicomponent fluids with non-random interactions. Molecular interactions are derived from the underlying design strategy (random, blocky, or sequence-based) and resultant interaction-matrix is used to assay phase behavior through complementary approaches (phase-field simulations, numerical free energy (NF) optimization based on [18], and eigen-mode counting from the Hessian matrix of the free-energy). See SI for details of approaches.

We investigate phase behavior through orthogonal approaches (SI Simulations; Figure 1). Briefly, we simulate the spatiotemporal evolution of volume fractions 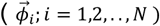 on a 2D grid employing conserved Model B dynamics. These physically-grounded, albeit computationally intensive, phase-field simulations (PF), predict spatial dynamics and steady-state properties. We then use an orthogonal numerical free-energy (NF) optimization, which is driven by non-physical mean-field dynamics [18] but can identify approximate number and composition of thermodynamic equilibria and is computationally faster (~1000x faster than PF). For both numerical approaches, we average over multiple initial conditions (concentrations, interactions) to measure *statistical* features of multiphase coexistence i.e. average number and composition of coexisting phases. Building on [17], we derive another estimate for the number of steady-state phases from the eigen-spectra of the Hessian matrix (SI Theory) as 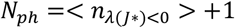.

We first these different approaches on a previously characterized model of multicomponent fluids with purely random interactions [17]. Here, eigen-mode counting can be approximated by an expression based on Random-Matrix Theory (RMT). Across a wide range of parameters, we find that simulations (PF and NF) and RMT agree on average number of phases at steady-state (Figures S1A-B), although NF mildly overestimates number of phases. Both simulations predict similar distributions of compositions of coexisting phases (Figure S1C). Finally, spatial dynamics from PF simulations show that random fluids exhibit the expected scaling laws (Figure S1D) that incorporate mechanisms of Ostwald Ripening and growth by coalescence [19]. Together, this suggests that the diverse methods we employ (eigen-mode, PF simulation, and NF) identify similar steady-state features for random fluids. To extensively characterize parameter space in what follows, we will especially leverage the eigen-counting method, which is ~10^6^ - 10^9^ faster than simulations.

## Blocky fluids

We first explore how structure in interactions influences phase behavior in a model of blocky fluids (Figure 2A). In this model, components (*N*) are split into blocks with strong and specific intra-block interactions (*m* = #_blocks, *ϵ_b_* =specific interactions). Crosstalk between species is introduced through random additive interactions (*ϵ_r_* ~ *N*(0, *σ*)). In the limit of very low cross-talk and strong interactions, it is straightforward to see that the number and composition of equilibrium phases matches individual blocks. Starting with two blocky phases (Figure 2B), we increase cross-talk between components by increasing *σ*. When cross-talk crosses a threshold, we find that the number of phases deviate from the block design (Figure 2B, simulations) and approaches expectations for a purely random fluid (Figure 2B; black line). While low cross-talk does not affect number of coexisting phases, we find that phase separation kinetics is modestly promoted (Figure S2A). We next sought to characterize how compositions of emergent phases change with increasing cross-talk. We measure the similarity of coexisting phases to the blocky phase (SI Analyses) through a relative angle *θ* (Figure 2C). Smaller values represent phases whose compositions overlap with blocks and larger values are orthogonal (and thus decoherent) to blocks. We find that the composition of coexisting phases becomes increasingly random beyond a cross-talk threshold (Figure 2C). These tradeoffs in number and composition are recapitulated by all our methods (Figures S2B-C).

**Figure 2:**
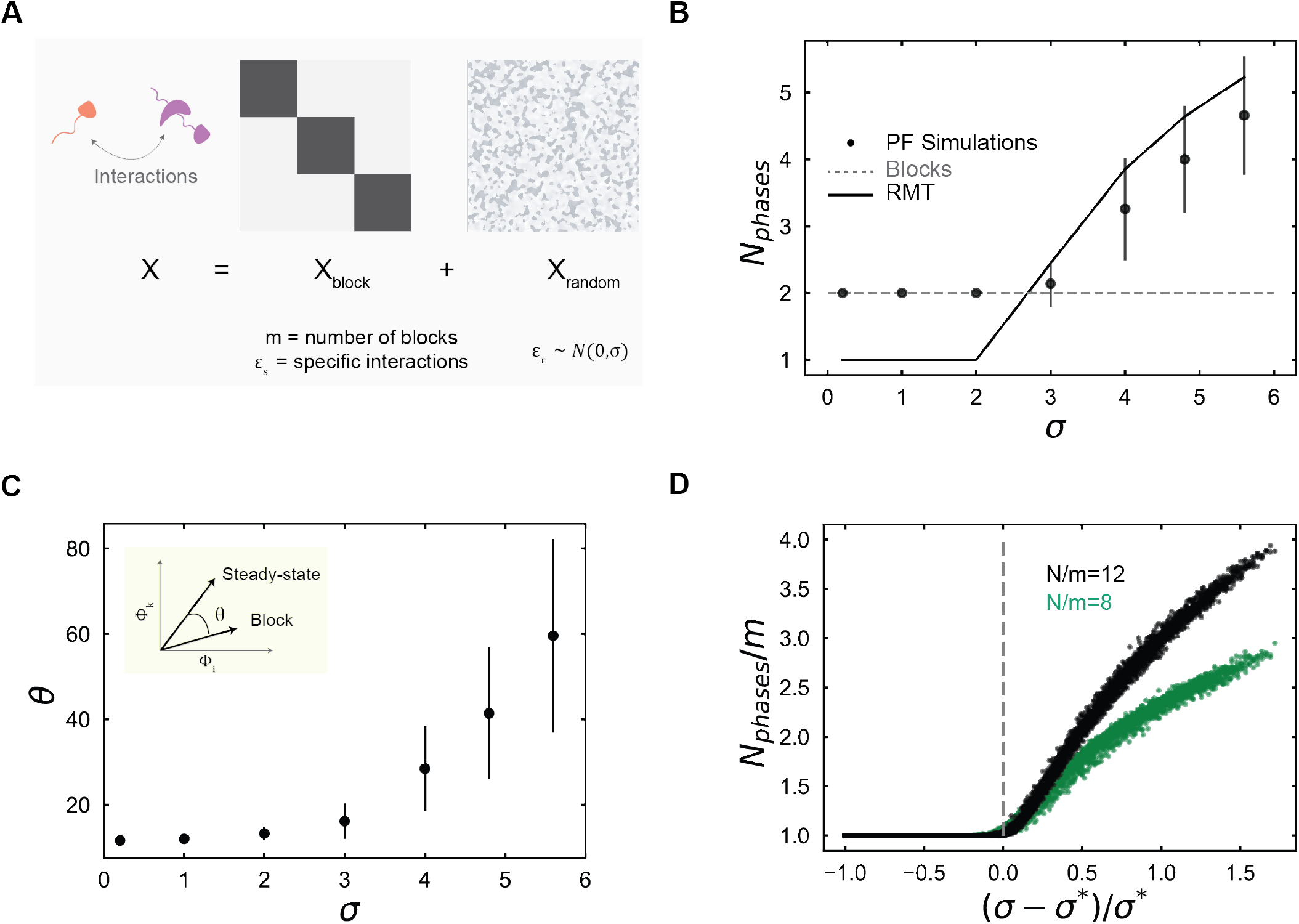
Specificity vs. cross-talk tradeoff in blocky fluids. **A.** A model of interactions in blocky fluids comprising a specific intra-block component and random additive crosstalk. **B-C.** Variation in number of steady-state phases and compositional similarity to a blocky phase with increasing cross-talk (*σ*). Black dots represent averages and dashes represent standard-deviation around averages over 50 PF simulations. In panel B, grey dashed line represents expectation from block considerations i.e. 2 specific phases, and black line represents theoretical predictions. All conditions are at N=16, m=2. **D.** A master curve describes scaled number of coexisting phases (*N_ph_/m*) versus renormalized crosstalk ((*σ* - *σ*\*)/*σ*\*) for specific values of 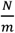 (black-=12, green=8). As specified elsewhere, *N* is number of species, *m* is number of blocks, *σ* is scale of cross-talk, and *σ*\*(*m, ϵ_ij_*) is inferred threshold.

We next sought to extensively characterize this tradeoff by leveraging eigen-mode counting. To ensure blocky phases in the absence of cross-talk, we intialize with a range of strong specific interactions (Fig S2D). Across many parameters, we measure the threshold of crosstalk required to disrupt blocky phases (*σ*\*). We find that crosstalk threshold, or *σ*\*, is largely independent of number of species (Figure S3A) but depends on number of blocks (m) and strength of interactions *ϵ_ij_* (Figure S3B-C). This dependence is well explained by a linear fit (Figure S3D). When we rescale crosstalk with our inferred fit, we find a universal collapse in the number of phases (Figure 2D). Here, the number of phases is normalized by the number of blocks, and is 1 below the threshold. Across a wide range of *N, m, ϵ_ij_*, conditions with similar *N/m* collapse on a master curve (Figure 2D – black and green points) but don’t depend on number of blocks (m) or strength of interaction *ϵ_ij_*. (Figures S3E-F). Overall, our results show a universal trade-off from specific phases to random decoherent phases for blocky fluids beyond a single rescaled threshold of crosstalk.

## Sequence-model

Towards more realistic models of biological molecules, we next consider interactions between species mediated by an underlying sequence. Motivated by a recent model [16], we describe each molecule as a sequence of length *R* (or *R* domains) with a continuous value associated with each position/domain. These values might describe protein features such as hydrophobicity or charge density or represent complementarity of DNA barcodes. For simplicity, we assume interactions between pairs of molecules is additive (Figure 3A, SI Theory), with interaction strength set by *J*. In our simulations molecules with similar sequences interact favorably (*J* < 0). Finally, we generate *N* species by normally sampling each feature as depicted in Figure 3A. This is akin to sampling a distribution of DNA barcodes where complementarity of interacting patches on the barcode is randomly sampled.

**Figure 3:**
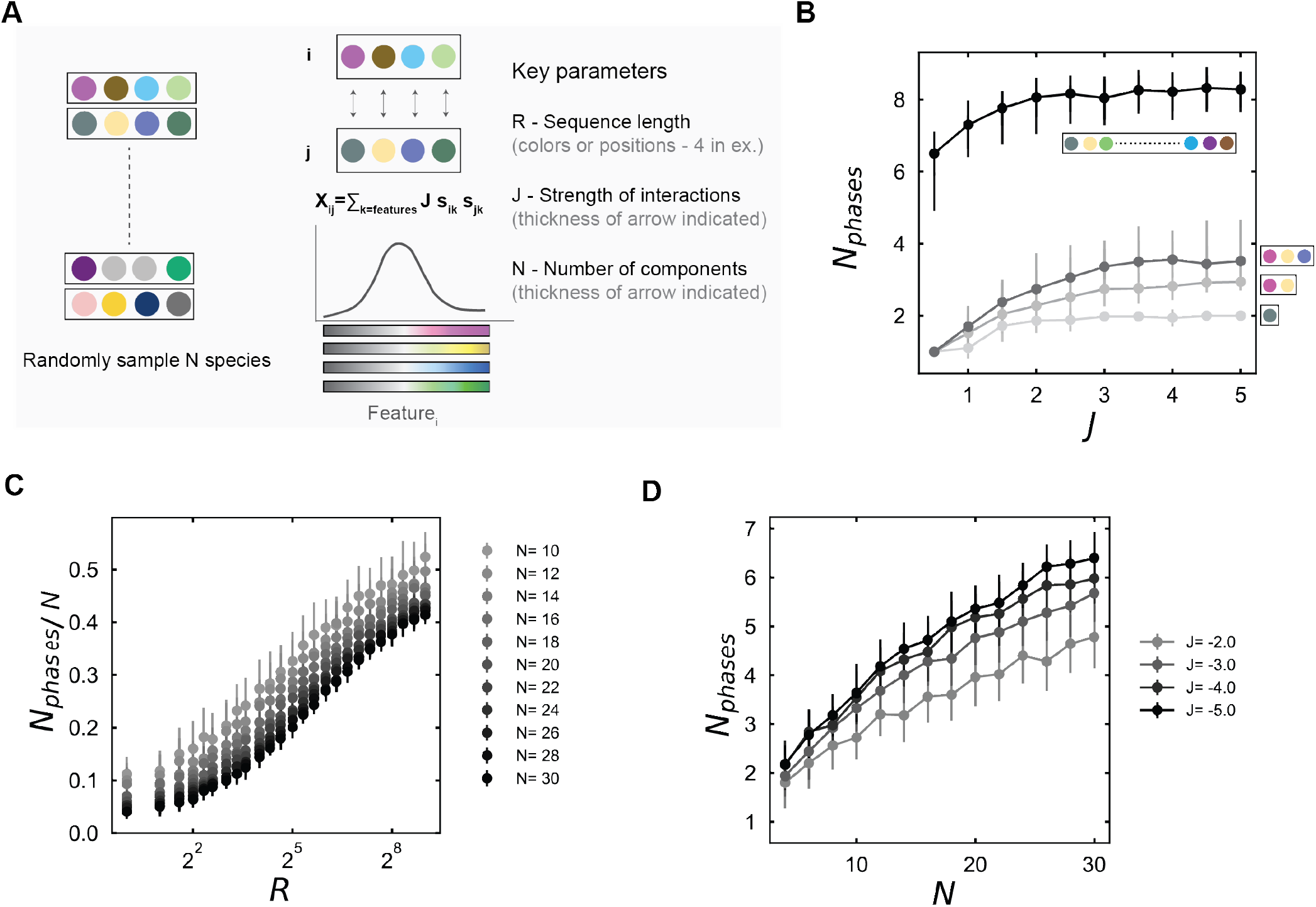
Sequence features constrain multiphase capacity. **A.** In the sequence-based model, each molecule contains a sequence of length R (or R interaction domains/features/patches). Each patch interacts with strength J to the same patch on another species. For generating an ensemble of N sequences, each feature is randomly drawn from a normal distribution. Interactions between two distinct species are additive across the features. **B.** Variation of number of steady-state phases with increasing strength of interactions for different sequence lengths. Grey lines represent R=1,2,3 and black line is R=512. In all conditions, N=16. **C.** Variation of scaled phase capacity (*N_ph_/N*) versus sequence length (R) for different number of species. Interaction strength is fixed at J=-2. **D.** Variation of number of phases versus number of species in the limit of strong energies. Here sequence length is fixed at *R* = 8. In all cases, markers represent average over 50 trials and dashes represent standard deviation over 50 trials.

To investigate how sequence encodes for phase behavior, we varied strength and number of features. Predictions from PF Simulations, NF, and eigen-mode counting agree on the number of steady-state phases with changing sequence length or interaction strength (Figures S4A-B). PF simulations show that when sequences are short (*R* =2), the maximum number of phases is ~ *R* + 1. This upper limit on phase capacity is validated by eigen-counting across a wider range of sequence features (Figure 3B; grey lines; Figure S5) when interactions are strong and sequences are short (*R ≪ N*). With larger lengths, sequences, and thus, interactions between molecules become less correlated. In the limit of large sequences (*R » N*), we find that the maximum phase capacity is limited by the number of species/barcodes (N) as 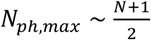 (Figure 3B; black line). Note that this limit collapses to the limit expected from purely random fluids that we showed before [17]. When number of species (N) or interaction strength (J) is constant, we find a logarithmic scaling between phase capacity and sequence length (Figure S4C-D; *N_ph,ss_* ∞ *γ* log(*R*) + *g*(*N,J*). This suggests that increasing sequence length provides only a modest increase in encoding phase capacity. The slope i.e. *γ*, is nearly independent of interaction strength but depends linearly on number of sampled species (Figure S4E). The intercept i.e. *g*(*J, N*), although non-monotonic, varies slowly at higher strengths (J) or sequence number (N) (Figure S4F). Based on this, we observe an approximate scaling collapse when interactions are not weak of the form 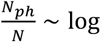 (*R*) across a range of conditions (Figure 3C). When length is fixed, our model predicts a linear increase in phase capacity with number of species (*N_ph_* α *Nlog*(*R*), which we observe across a range of interactions (Figure 3D). This shows that even generating random sequences gives rise to a linear increase in phase capacity. Together, our results show that sequence features constrain multiphase capacity in characteristic ways.

In summary, we explore how non-random molecular interactions map to spontaneous phase behavior. Towards this, we employ different methods, namely phase-field simulations, mean-field NF optimization, and eigen-mode counting, which we validate for a previously characterized model of random fluids (Figure S1) [17]. We first study a model of blocky fluids, where interactions between components comprise both specific intra-block and random crosstalk parts. We identify a universal trade-off that leads to a cross-over from block-specific phases to random decoherent phases beyond a single rescaled threshold (Figure 2), suggesting interesting but distinct parallels to tradeoffs in Hopfield networks and molecular self-assembly [20]. We next define and characterize a sequence-based model of interactions (Figure 3). When sequences are randomly generated, we show that phase capacity grows logarithmically/linearly with sequence length (R) / number (N). We find that the maximum phase capacity is limited by sequence length or number of species i.e., min 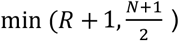. Together, these results connect features of interaction networks to emergent multiphase coexistence.

Programmable DNA barcodes or charged peptides [21,22] offer experimentally instantiable avenues to test our predictions. While we employ an additive model for simplicity, the link between sequence and interactions is often non-linear [23] and will be an important avenue for future research. More generally, developing theories and methods to predict steady-state coexistence and nucleation kinetics in multicomponent and multiphasic fluids represents an exciting frontier for complex living materials.

## Acknowledgements

K.S was supported by the NSF-Simons Center for Mathematical and Statistical Analysis of Biology at Harvard (award number #1764269) and the Harvard FAS Quantitative Biology Initiative. M.P.B was supported by ONR N00014-17-1-3029 and the Simons Foundation.

## Author contributions

K.S. and M.P.B conceived the project. K.S. developed simulations and analyses, and wrote a first draft of the paper. K.S. and M.P.B edited and revised the manuscript.

## Materials and Methods

Custom code was written in python for phase-field simulations, suitably modifying interactions to non-random models from https://github.com/krishna-shrinivas/2021_Shrinivas_Brenner_random_multiphase_fluids. NF approach was adapted from [18]. More details are provided in Supplementary Information.

---

**Figure S1:**
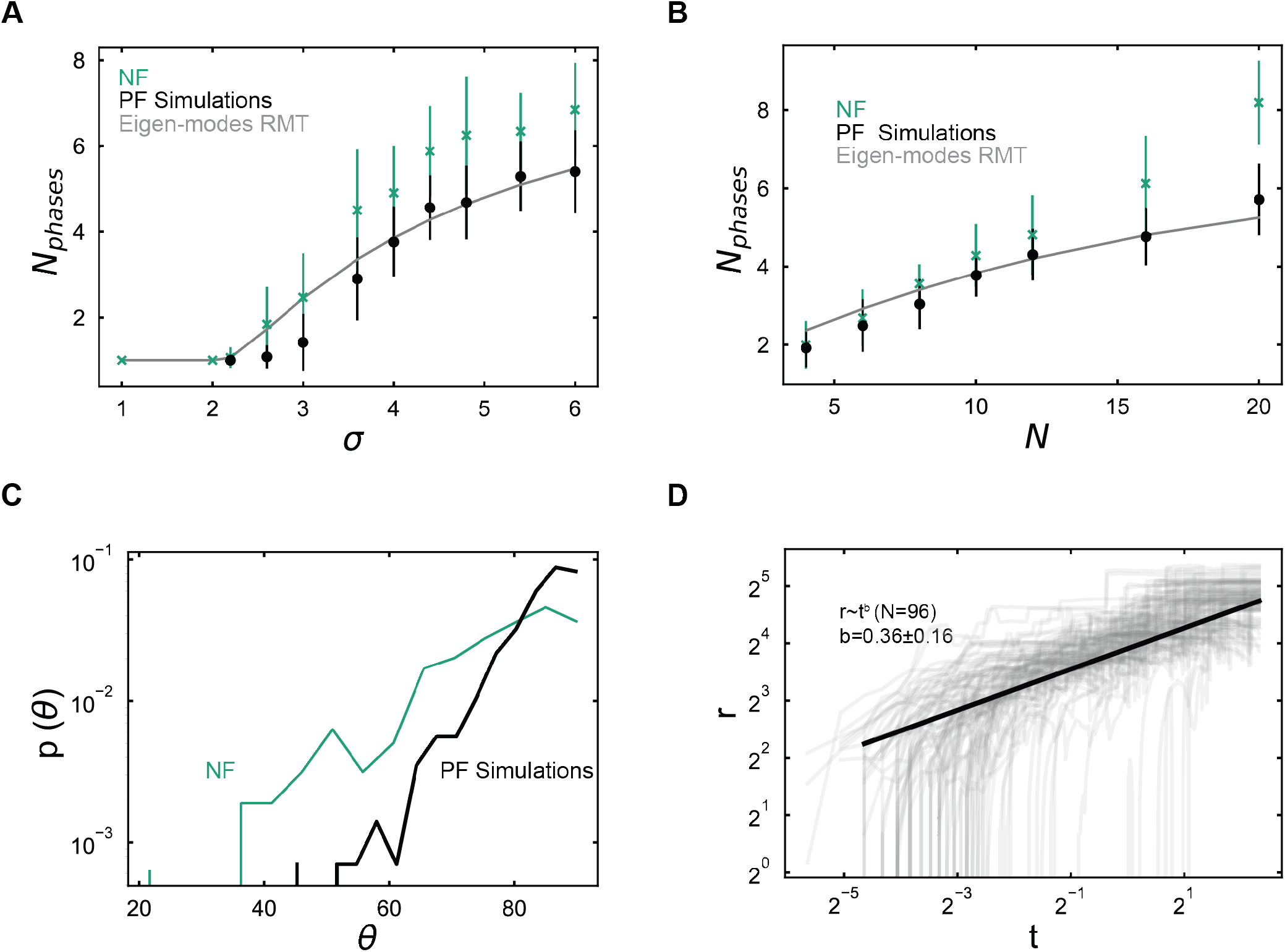
Dynamics and multiphase behavior in random fluids. **(A-B)**. Variation of number of steady-state phases versus width of interaction distribution and number of components. Green dots/dashes are means/standard-deviations over 50 NF trajectories and black dots/dashes are means/standard-deviations over 50 PF Simulations. Grey lines are derived from random-matrix theory based approximation of eigen-mode counting. In panel A, the number of components if fixed at N=16, and in panel B, the width of interactions is *σ* = 5. **(C)** Distribution of angles between compositions (*θ*) of coexisting steady state phases from PF simulations (black) and NF model (green). **(D)** Statistics of merger/growth events of phases < *r* > versus time (t), with an inferred power law scaling exponent *b*, for N=16 species and *σ* = 4.8.

**Figure S2:**
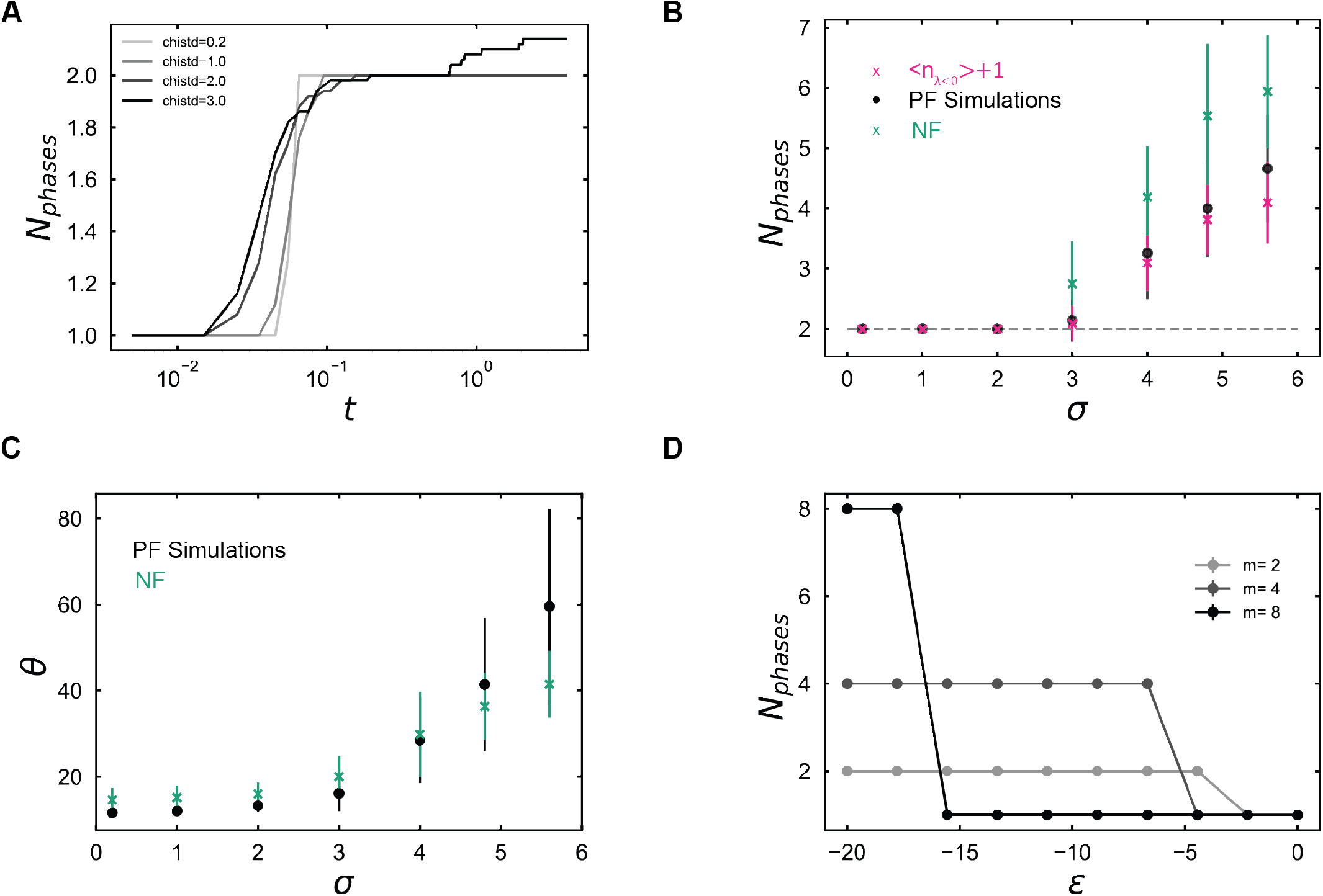
Dynamics and multiphase behavior in blocky fluids. **(A)** Number of phases versus simulation time for increasing crosstalk (darker colors) – lines represent average over 50 phase field simulations. **(B)**. Variation of number of steady-state phases versus crosstalk by phase-field simulations (black), NF (green), and eigen-mode counting (pink). Grey line represents number of blocks/phases. **(C)**. Variation of relative composition of steady-state phases versus specific blocky phase with increasing crosstalk by PF simulations (black) and NF (green). **(D)** Variation of number of steady-state phases versus strength of intra-block interactions for increasing number of blocks (m=2, m=4, m=8) in the limit of very low cross-talk for N=16 components. In panels with markers, symbols represent average over 50 trials and dashes represent standard deviation over 50 trials.

**Figure S3:**
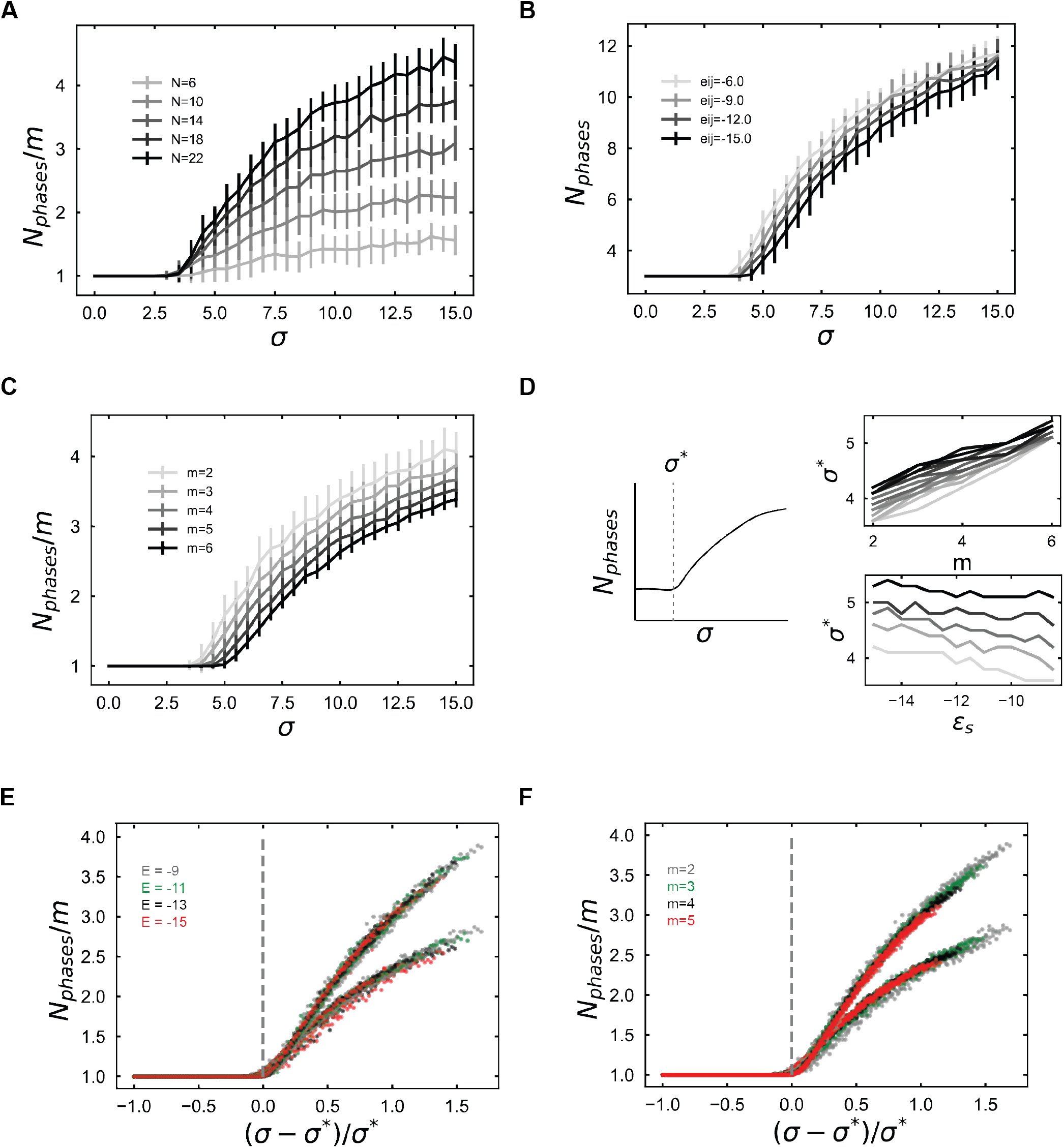
Specificity and cross-talk in blocky fluids. **(A-C)**. Specificity-crosstalk trade-off in number of phases with increasing random crosstalk (*σ*) on x-axis and increasing number of components (darker colors) for N=16 components and m=2 (S3A), increasing interaction strength for N=27 species and m=3 blocks (S3B), and increasing number of blocks for *ϵ* = −9. (S3C). In panels A,C, the number of phases is normalized by number of blocks. **(D)**. Linear variation of crosstalk threshold with number of blocks and energy of specific interactions. **(E-F)**. A master curve describes scaled number of coexisting phases (*N_ph_/m*) versus renormalized crosstalk ((*σ* - *σ*\*)/*σ*\*) where individual points are false colored by specific interactions (S3E) or number of blocks (S3F). In all panels, markers represent average over 50 trials and dashes represent standard deviation over 50 trials of eigen-mode based calculations.

**Figure S4:**
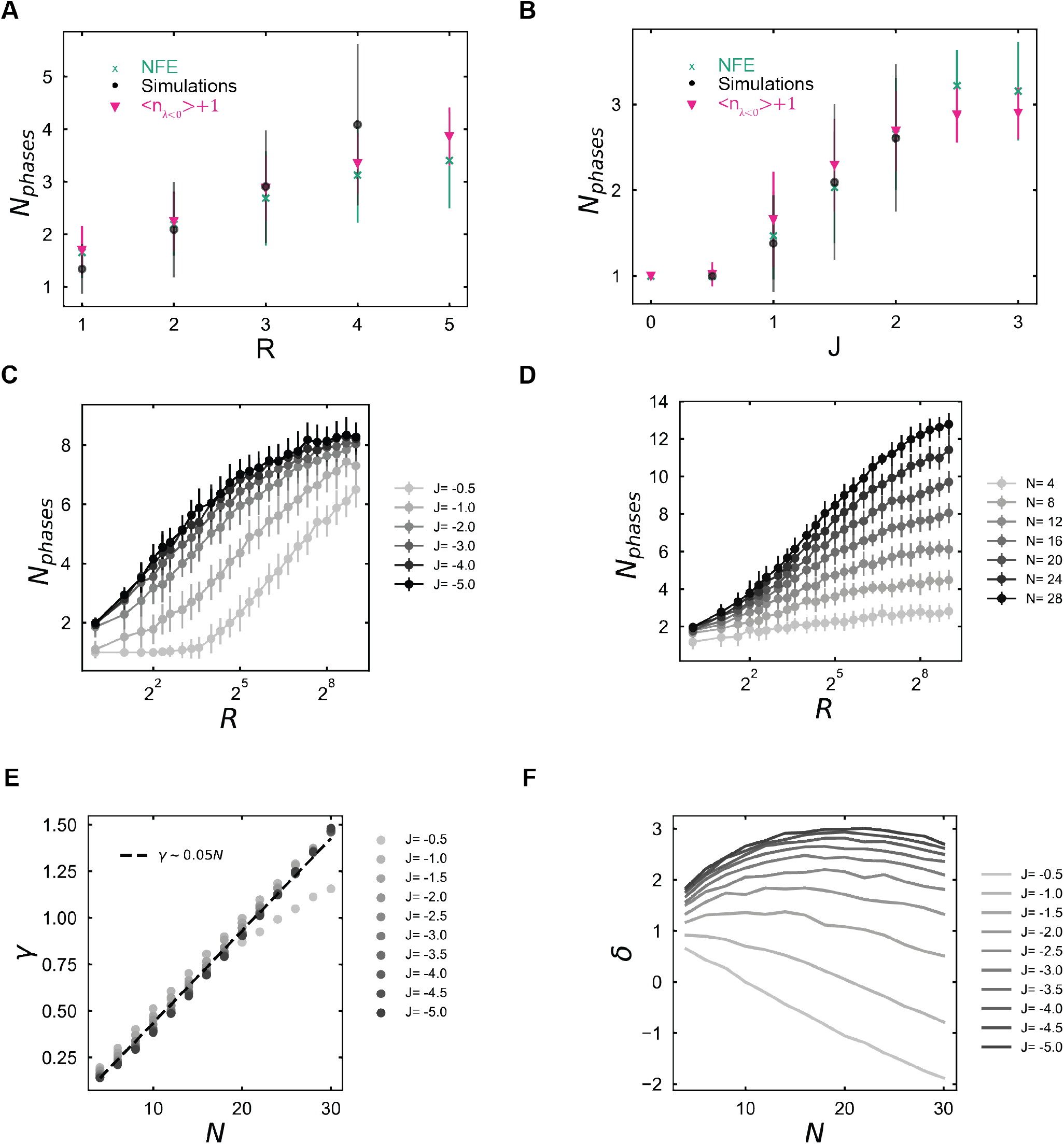
Scaling relationships in sequence-based model of interactions. **(A-B).** Variation of number of steady-state phases with sequence length R) and interaction strength (J) using phase-field simulations (black), NF (green), and y eigen-mode counting (pink), For panel B, J=-2, and for panel C, R=2, and in both conditions N=16 **(C-D)**. Variation of phase capacity with increasing strength of interactions J (S4C) or number of species (S4D) for different number of features (x-axis). Capacity is limited by total number of components N=16. **(E-F)** Variation of scaling coefficient slope (*γ*) and intercept (*g*) with N (x-axis) for different strengths of interactions (J, darker colors represent stronger interactions) derived from eigen-mode counting. Panel A shows a linear dependence of the slope on N for all energies. In all panels, markers represent average over 50 trials and dashes represent standard deviation over 50 trials of eigen-mode based calculations.

**Figure S5:**
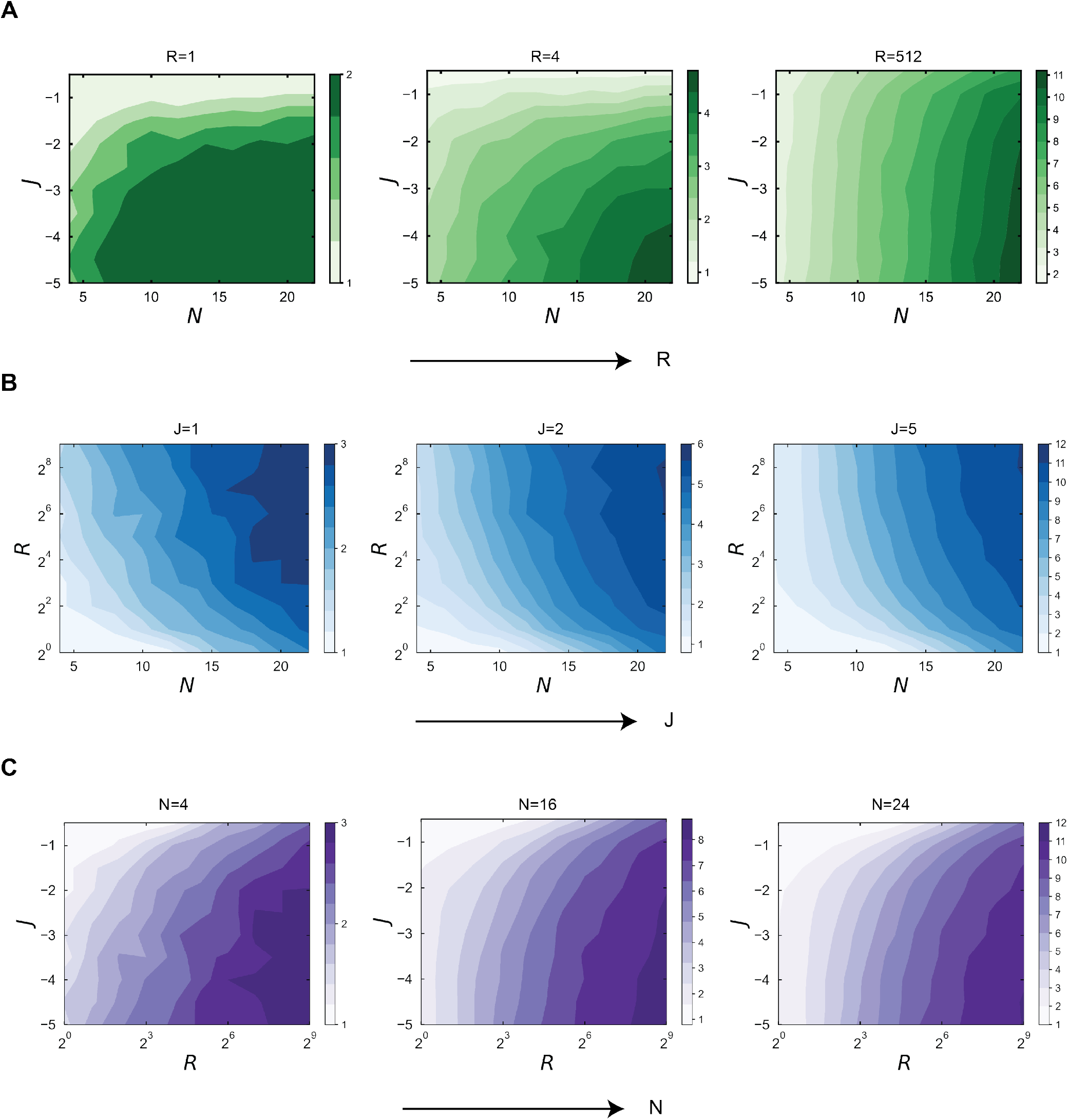
Sequence-features constrain number of encoded phases. **(A-C)**. Contour plots of mean number of phases for all key parameters of the sequence model i.e, sequence length/features (R), interaction strength (J), and number of sampled species (N). Each row represents a slice of the 3D contour with (A) J vs N, (B) R vs N and (C) J vs R. In each row, the third axis increases from left to right. In all panels, mean phases are computed over 50 trials of eigen-mode based calculations. The supplementary information for this manuscript is arranged into separate sections for theory, simulation, and analyses. 5 supplementary figures are found above.

## Supplementary Information

### Theory

In this paper, the underlying thermodynamic model employed follows the mean-field regular solution/Flory-Huggins style approach. The free-energy is defined as:

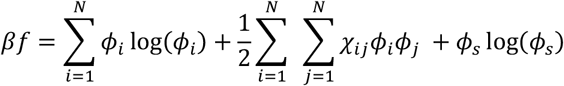

Where the *N* is the number of interacting species, *ϕ_i_* is the volume fraction of species *i, φ_s_* = 1 - ∑_*i*_*ϕ_i_* is the volume fraction of the solvent, and *χ_ij_* α *ϵ_ij_* is the Flory parameter that is proportional to interaction strength. We assume the solvent is inert, molecules do not self-interact and an equimolar solution i.e., 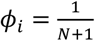. The Hessian, whose eigen-spectra are used to count unstable modes, is calculated as:

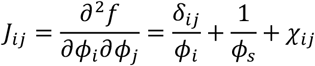

Interactions between species (*χ_ij_*) depends on ensemble and we outline briefly how they are specified.

#### Random fluids

For fluids with random interactions (Figures 1, S1), the interactions are drawn as i.i.d values from a gaussian distribution with zero mean i.e., *χ_ij_* ~ *N*(*μ* = 0, *σ*).

#### Blocky fluids

Interactions between components comprise a specific and random noisy component (Figures 2, S2, S3). Species (N) are split into m blocks/groups with the following interaction energies:

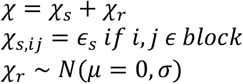

#### Sequence-based fluids

Interactions between components arise from additive interactions from underlying sequence of species (Figures 3, S4, S5). Briefly, each species has *R* features or sequence locations, whose values *s_ik_* are independently drawn from a normal distribution for each feature/position. A total of N species are generated per trial through this procedure. Here the subscript index *i* (1,…,*N*) denotes species and *k* denotes feature/position within species (1,…,*R*). The pairwise interactions between distinct species *i* and *j* is then defined:

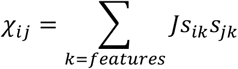

### Simulations

Simulation techniques employed in this manuscript are described below. More generally, in each simulation techniques, multiple trials or trajectories were performed with similar parameters – randomly generating a concentration and interaction matrix/ensemble for each trial. Unless otherwise stated, the reported results are typically averaged over 50 simulations.

#### Phase-field simulations

The spatiotemporal volume fractions of N species *φ_j_*(*r, t*) evolve according to the multicomponent Model B equations, previously introduced in [17]. Briefly, the equations are:

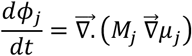

Where the mobility used is *M_j_* = *Mϕ_j_* recapitulates Fickian diffusion in limit of dilute inert solvent. The chemical potential term includes an interfacial stabilization term (*κ* ∝ surface-tension) and is described as:

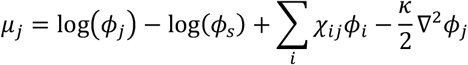

All simulations are performed on a discretized 2D mesh, grid size 64 × 64 with a timestep of *dt* = 2*e* - 6 and simulations are run for atleast 25 × 10^6^ timesteps to ensure convergence. The numerical scheme deploys an implicit-explicit description, previously described I in [17] with a fourth-order stabilization term included for convergence. The initial concentrations are sampled as equimolar compositions with uniform noise added of magnitude 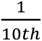 of absolute value. For each individual trajectory, a composition and interaction matrix are sampled.

#### Numerical free-energy (NFE) optimization

We leverage a recently developed method, described in detail in [18], to identify the free-energy by numerical optimization of coupled ODEs. Briefly, the NFE method leverages linear irreversible thermodynamics to set up relaxation dynamics for the compositions of species so that the solution of equations converge to minima that have identical chemical potentials and mechanical pressures. The governing equations are:

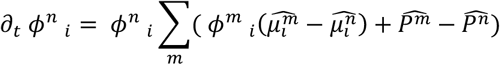

Where 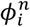 represents volume fraction of species i in phase n, 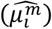 is the non-dimensional chemical potential of species *i* in phase m, and 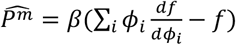 is the non-dimensional mechanical pressure of phase *m*. Together, the steady-state of these equations converge to states that are in the neighborhood of equilibrium coexisting phases. The numerical implementation provided in [18] is directly employed, except the interaction matrices are generated as previously described.

For running 50 trials of a particular set of parameters (block model) the approximate times for convergence are identified on a single node.

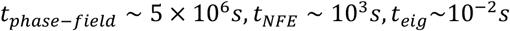

### Analyses

For each of the simulations, the final outcomes are a set of *k* phases with compositions 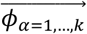, identified either from mesh data following PCA and clustering as in [17] or from clustering following [18].

*Analyses on compositional similarity* between two phases *α,β* is measured by the angles between the two vectors 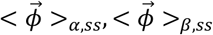. Angles close to 90’ represent orthogonal phases whereas angles close to 0 represent largely similar phases by composition. For Figure 2, the reference phase is used as one of the blocky phases, highly enriched in all block components but lacking non-block species. Probability distributions in Figure S1 are computed by identifiying distribution of angles between multiple coexisting phases over 50 trajectories.

#### Merger statistics

In Figure S1C, the phase-field simulation dynamics data is taken and distribution of phase sizes over time is identified. By masking the mesh data with phase labels, the average size of individual droplets is measured over time. This data is collected across 50 trajectories, and the mean-increase in size is fit to a power of the form < *r* >~ *t^b^*. The inset text reports the average as well as the standard deviation in growth of droplet sizes.

#### Scaling laws

For the scaling laws identified in Figure 3, Figures S4–S5, the eigen-modes are computed a wide range of number of species N, features R, and strengths J across 50 random samples per parameter condition. For a given (N,J), the variation in mean predicted phases (< *N_ph_* >) is linearly regressed against log(fi) to identify a slope *γ* and intercept *g*(*N,J*), plotted in Figures S5A-B. For all conditions, *N_ph_vs* log(*R*) fit the data well, with regressed *R*^2^ > 0.99.

#### Threshold-fit

For blocky fluids, we identify the threshold for crosstalk (*σ*\*) as the minimum value beyond which the average number of phases deviates more than 5% above the number of blocks. We identify the thresholds across a wide-range of thresholds and identify a linear fit of the form (Fig S3D) below:

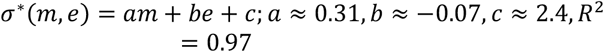

Here, m is the number of blocks and *ϵ* is strength of specific interactions.

